# Amyloid Folding: an Origami-Based Approach

**DOI:** 10.1101/2025.06.12.659339

**Authors:** Luís Maurício T. R. Lima

**Author notes:** To whom correspondence should be addressed*: Luís Maurício T. R. Lima – Faculdade de Farmácia, Universidade Federal do Rio de Janeiro – UFRJ, CCS, Bss24, Ilha do Fundão, 21941-902, Rio de Janeiro, RJ, Brazil. **Author information** Luís Maurício T. R. Lima.

## Abstract

Amyloid is an ordered folding pattern that involves a cross-beta arrangement, with hydrogen-bonding between adjacent chains in the fiber elongation axis, and side-chain interactions perpendicular to it, forming steric and/or polar zippers intra- and/or inter-chains. The arrangement of the polypeptide backbone and the side-chain interactions can be arranged with different morphologies that are repeated along the fiber growth axis as stacking units.

A simple representation of amyloid folding of proteins using paper origami is reported here, which can be used to visualize the β-sheet along fiber axis and the perpendicular variability in topological arrangement. The model uses regular office paper, in the form of multiple linearized polypeptide chains, arranged in parallel or anti-parallel forms, connected by hydrogen bonds. Alternating mountain/valley creases along the growth axis of the fibrils and, over the front/back side-chains, results in a pleated sheet that can be used to study the topological arrangement of amyloid fibrils and the polar/steric zippers. The present models can be used as teaching tools for understanding the structural and molecular basis of amyloid folding, chain growth, homo- and cross-seeding, and other features of amyloid function in health, disease and biotechnology.

**Graphical Abstract:** 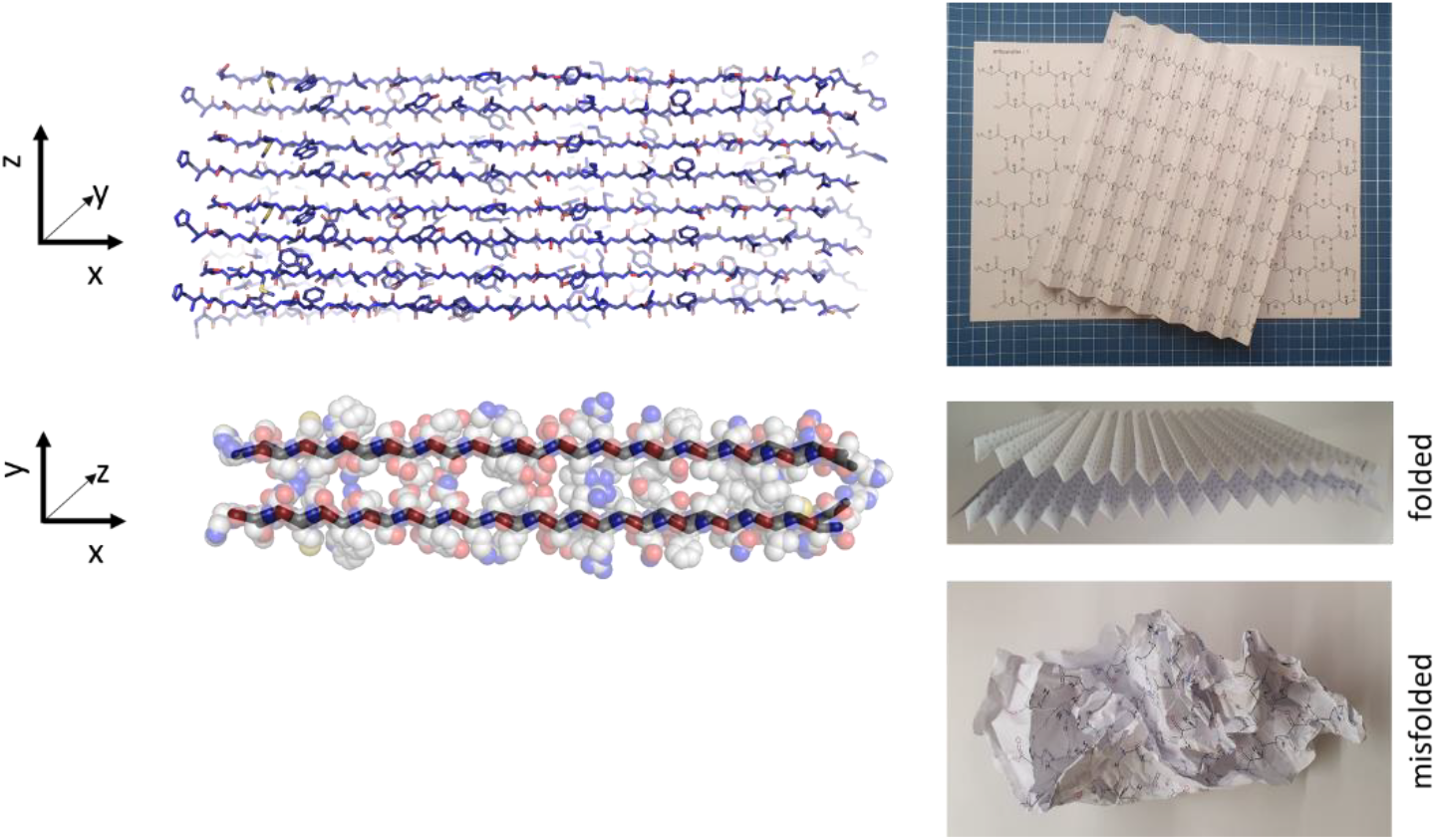

**Key features:**

- Introduces a **physical origami model** of amyloid β-sheet structure using paper folding.
- Demonstrates how **hydrogen bonding and side-chain interactions** can be conceptualized in 3D.
- Models **morphological polymorphism** and **parallel/antiparallel alignment**.
- Useful for **teaching complex biophysical concepts** like cross-seeding, fibril growth, and molecular stability in amyloids.
- Highlights the **educational value** for biochemistry, structural biology, biomedical and material sciences.

## 1. Introduction

In his groundbreaking book “*On Growth and Form”* (Thompson *et al*. 1942), the biologist and mathematician Sir **D’Arcy Wentworth Thompson** recalled the saying that:

> “*chemistry would only reach the rank of science, in the high and strict sense, when it should be found possible to explain chemical reactions in the light of their causal relation to the velocities, tensions and conditions of equilibrium of the component molecules*”.

Sir Thompson sought to emphasize the physics and mechanics of the **form** and structure of biological entities in his book. While he focused on living organisms, by the time his book was published (1^st^ edition 1917, 2^nd^ edition 1942) the foundations for the understanding the structure and function of atoms, molecules and macromolecules has already been laid.

In the 1930’s, Prof. **William Thomas Astbury**, at the Leeds University, studied fibers of different origins, and observed that proteins transformed from, what he called, an α-to a β-form, the later showing x-ray diffraction spacing of 4.65 Å. Prof. Astbury proposed a model in which this distance represents the spacing between the main chains of polypeptide chains, hold together by hydrogen bonds connecting amide C=O and N-H from adjacent chains (Astbury 1933) ^1^ (**Fig. 1**).

**Figure 1.**
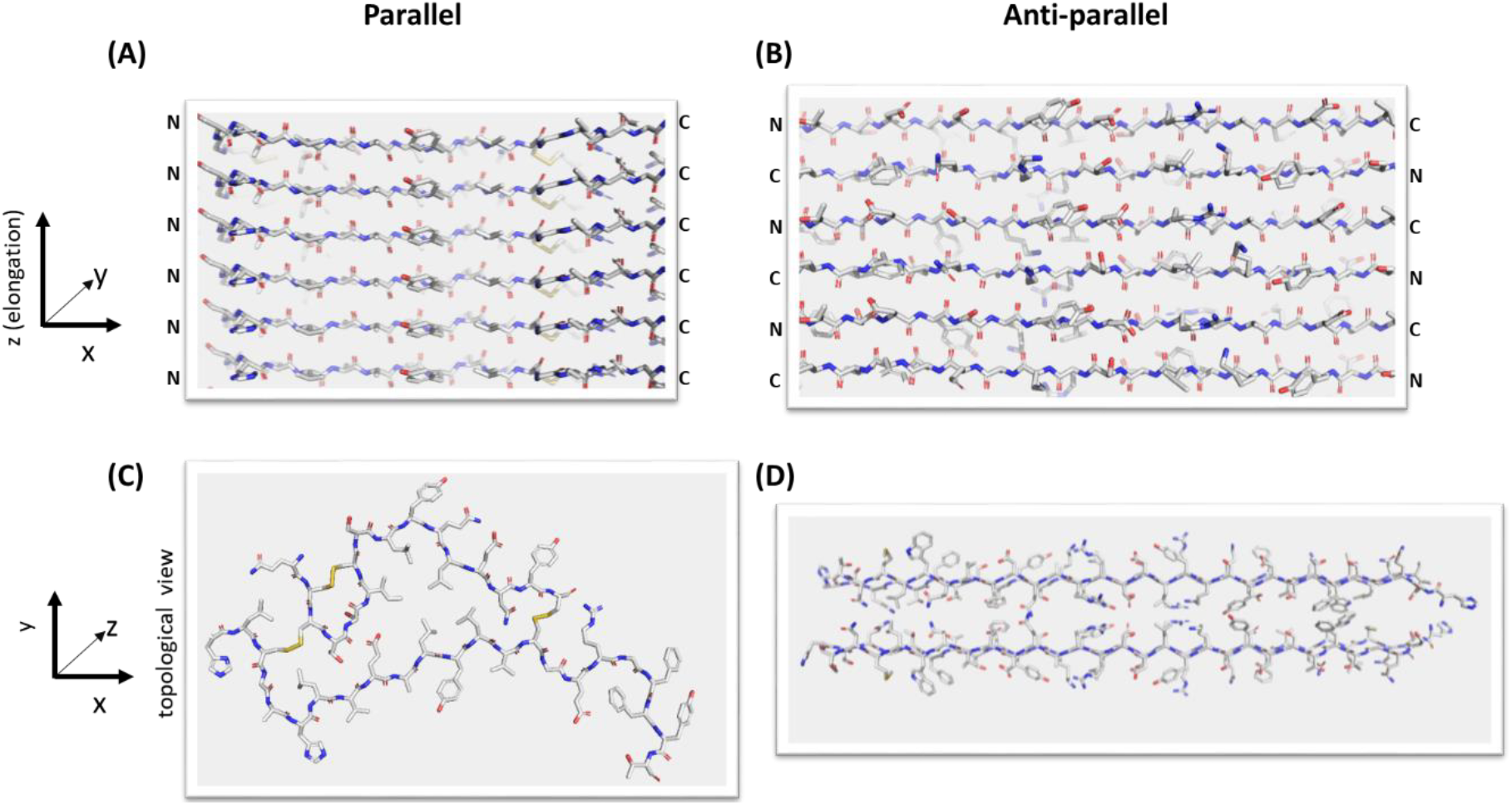
Amyloid folding. Representation of parallel (**A** and **C**; insulin; PDB ID: 8SBD; cryoEM) and anti-parallel (**B** and **D**; glucagon; PDB ID: 6NZN; NMR) amyloid folding. Two orientations: front view of the amyloid elongation axis depicting the stacks of polypeptide chains layers (**A** and **B**), and top view (perpendicular to previous one) evidencing the topological arrangement. Not to scale between panels.

Drawings and physical models were also important for understanding of the structure of DNA (Watson & Crick 1953) and globular proteins (Perutz *et al*. 1960; Kendrew 1961), leading to Nobel Prizes in Medicine/Physiology (to Prof. **James Watson**, Prof. **Francis Crick** and Prof. **Maurice Wilkins**) and in Chemistry (to Prof. **Max Perutz** and Prof. **John Kendrew**) in 1962. Nowadays, physical models and computerized virtual models are widely used to represent chemical structures and can support both scientific research and teaching.

Large globular protein molecules are difficult to represent with all-atoms models. The alternative cartoon drawing of the main secondary structures of the polypeptide backbone are routinely used in both programs (e.g.: <u>ChimeraX</u>, <u>PyMOL</u>, <u>MolMol</u>, embeded tools in RCSB and <u>UNIPROT</u>), toolkits (Garratt *et al*. 2015), 3D-printed models (De Baere *et al*. 1992; Kerwin 2019; Popa & Saitis 2022), and origami (Demaine & O’Rourke 2007; Reißer *et al*. 2018; Azulay *et al*. 2020, 2022; “Molecular Origami: Build 3D models of PDB Structures”; “Molecular Origami: Build 3D Paper Models of Protein Domains”). However, the chemical basis for amyloid folding and topological arrangement are not easily understood. While solving the paradox of protein folding from Prof. Cyrus Levinthal (Levinthal 1969), on “how to fold graciously”, has awared the 2024 Nobel Laureates in Chemistry to Profs. **David Baker**, Prof. **Demis Hassabis** and Prof. **John Jumper**, the interplay between complexity and simplicity of the polymorphic nature of protein folding remain unsolved.

Amyloid is a fibrillar morphology of proteins, generally referred to as **misfolding** compared to said native structure. More than 50 endogenous or iatrogenic diseases are have been linked to amyloid (Buxbaum *et al*. 2022), although amyloid can also be formed by proteins unrelated to diseases (Chiti *et al*. 1999). The Nobel Prize of Physiology/Medicine was also awarded to Daniel Carleton Gajdusek (1976) and Stanley B. Prusiner (1997) for their research in amyloid and diseases, although no amyloid structure had yet been solved at the time. Today, more than 500 amyloid structures have been solved by cryoelectron microscopy (cryoEM) or solid-state Nuclear Magnetic Resonance (ssNMR), enabling the understanding of key chemical and structural properties, such as:

- hydrogen-bonding between amide groups of adjacent **parallel and/or antiparallel** chains along the fibril elongation axis forming a **β-sheet**, with average distance of **4.7 Å**;
- side-chains interactions inter- and intra-chains in the same or adjacent topological “planes” perpendicular to the elongation axis, characterizing the **cross-β** pattern in amyloid fibrils.

The interactions between the side chains can vary in nature, forming electrostatic or other polar interactions (Perutz *et al*. 1993; Perutz 1994) (denominated **polar zippers**) and hydrophobic effects and packing restricted by van-der-Waals volume of the side chains (Ivanova *et al*. 2009) forming **steric-zippers**, by analogy to leucine zippers (Landschulz *et al*. 1988; Meyer 2015).

Since the fibril resembles to a stacking of repeating topological planes (or layers) connected by hydrogen-bonding between amide groups with contributions from side-chains within and between these layers, the structure of the amyloid folding can be represented in a simplified form by showing the geometric arrangement of the chains in such layers, as elegantly proposed by Prof. **Michael R. Sawaya** and Prof. **David S. Eisenberg** (Sawaya *et al*. 2021), and available in the **Amyloid Atlas** (https://people.mbi.ucla.edu/sawaya/amyloidatlas/). Still, understanding amyloid structure can be complex depending on individual ability to visualize polypeptide chains orientations, the “zipper” concept and the fibril elongation axis. To address this issue and introduce the educational and scientific community to the amyloid concept and science, a simple origami model for amyloid folding is proposed here.

## 2. Experimental

Polypeptides chains composed of L-amino-acids were designed using the free *ChemSketch* software (ver. 2024.2.0; https://www.acdlabs.com/resources/free-chemistry-software-apps/chemsketch-freeware/). Other alternatives to *ChemSketch* (https://alternativeto.net/software/acd-chemsketch/) may work for modeling, although they have not been tested for the purpose of this work. A polypeptide chain was represented as a linear structure aligned in one direction of the page. Additional chains were replicated and distributed along the growing fiber axis to simulate adjacent strands within a β-sheet. Using the “*Flip Left to Right*” function, chains could be displayed in parallel or antiparallel orientations. Hydrogen bonding interactions between chains were represented by dotted lines and positioned to reflect close proximity between backbone atoms.

The printing process involves creating double-sided pages. Care must be taken to properly align the front and back sides, either by manual inversion or through automatic duplex printing (depending on the printer capabilities). Alignment accuracy should be verified by holding the printed page against a light source to ensure proper overlay of the chains on both sides. Improper alignment—such as printing the same side twice—will result in incorrect structural representations.

The *ChemSketch* file contains eight pages, in the following order:

1. 8 parallel chains, front
2. 8 parallel chains, back (flipped)
3. 8 anti-parallel chains, front
4. 8 anti-parallel chains, back (flipped)
5. 16 parallel chains, front
6. 16 parallel chains, back (flipped)
7. 16 anti-parallel chains, front
8. 16 anti-parallel chains, back (flipped)

Each front & back pair should be printed one in each side of the paper. Diagrams can be printed (or hand-draw) in varying paper sizes, such as office, letter, A4, A3, A2 or other according to user preferences and availability.

Other figures using high resolution structures with coordinates deposited in the RCSB were prepared using PyMOL (https://pymol.org/). Other alternative software (https://alternativeto.net/software/pymol/) can be used according to preferences, skills and availability to the same modelling strategy.

## 3. Results

Amyloid folding can be understood as arrays of polypeptide chains, parallel (**Fig. 1A and Fig. 1C**) or anti-parallel (**Fig. 1B and Fig. 1D**), with two main axes: the fibril growth / elongation axis (Z; **Fig. 1A and Fig. 1B**) and a perpendicular stack of planes (X-Y; **Fig. 1C and** **Fig**. 1D) with the topological distribution of the polypeptide chain. The elongation axis is characterized by hydrogen bonding between amide groups of adjacent chains from the stack, while the chain has the topological arrangement maintained by forces driven by side-chain interactions as polar and/or steric zippers and hydrophobic clusters (**Fig. 1C and Fig. 1D**).

Four groups of **parallel** and **anti-parallel** polypeptide chain arrays (8 or 16 chains), forward (N-to-C terminus) and backwards (C-to-N terminus), were generated for A4 office paper size, and can be satisfactorily printed in paper with basis weight of about 75 g/m^2^, but can be performed to other desired density and standard or custom paper size according to the capabilities of the program (**Fig. 2**). Higher density (120 g/m^2^) are not satisfactory for A4 to office paper size, although it may be adequate to larger paper sizes (e.g.: poster size, 70 cm x 120 cm). The models can be printed in papers of different colors (e.g.: blue, yellow, pink, green), in order to represent different fibrils, and mounted adjacent to each other. A single A4 paper sheet can accommodate 8 or 16 polypeptide chains (available in **Supporting Material**) with sufficient space to be handled manually and folded without much difficulty even for people little skilled in origami. The *ChemSketch* file contains models with eight parallel polypeptide chains contains the representation of 17 side chains, and models with 16 parallel polypeptide chains contains the representation of 33 side chains, requiring the same amount of alternating mountain/valley folds to complete the model. These models can be useful to model amyloid folding patterns with small or large polypeptide chains respectively. Longer or shorter polypeptide chains can be obtained by proper editing the *ChemSketh* file.

**Figure 2.**
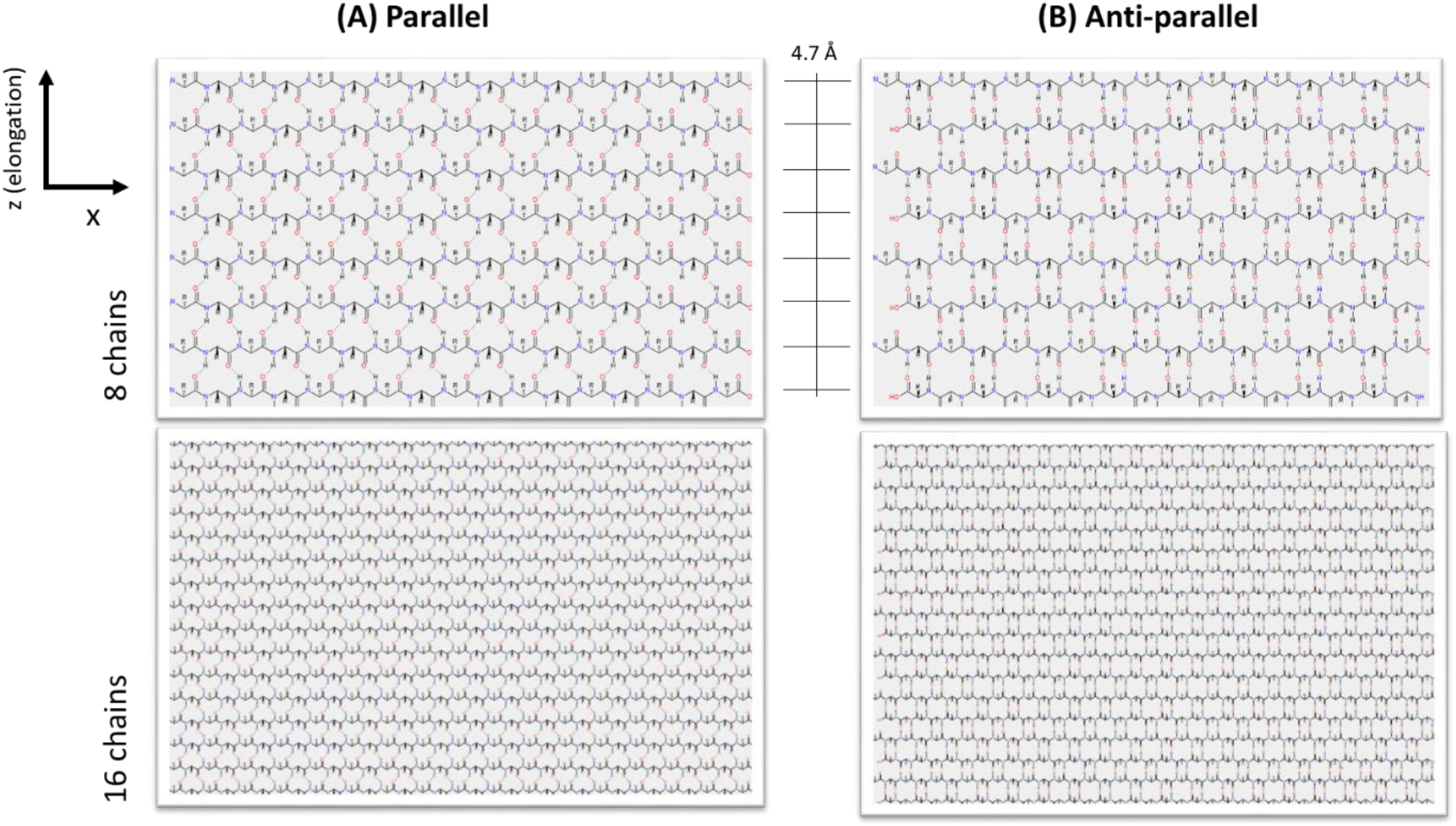
Polypeptide chain drawing. Representation of the printed form of the polypeptide chains for paper folding and study of amyloid folding. (**A**) Parallel chains; (**B**) Anti-parallel chains.

Folding direction should preferably follow the origami convention of **mountain** and **valley** fold, which are opposing each other. Mountain fold should be done over the R-group represented at the front of the paper plane (bold bond), connecting the corresponding R-group from adjacent chain, while valley fold should be done in opposite direction using the R-group represented to the back of the paper plane (dashed bond) (**Fig. 3**).

**Figure 3.**
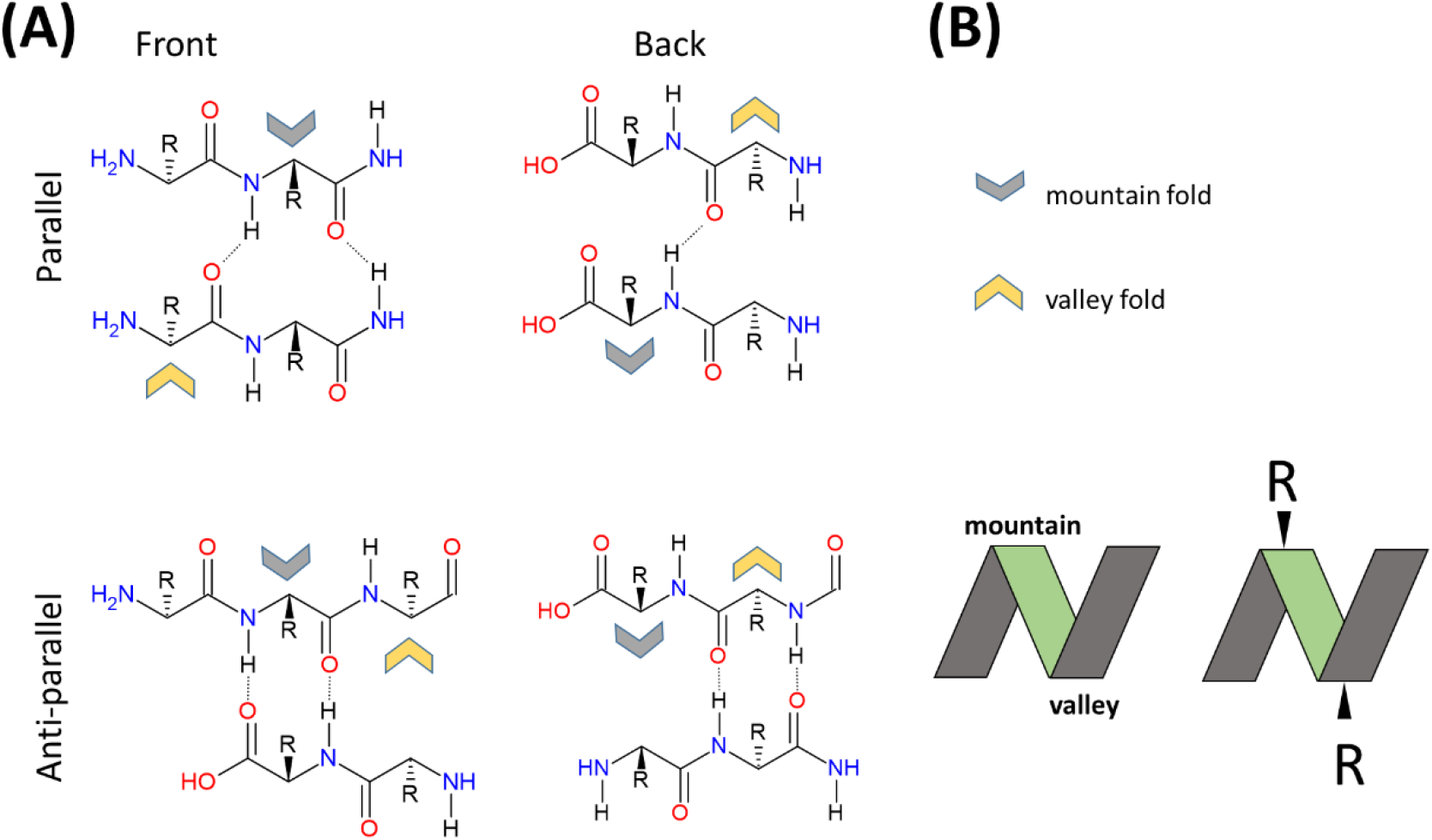
Detail of the chains drawings with print and folding direction. **A)** Fragment (chemically incomplete) of the polypeptides and the chain direction (N-to-C: forward and backward). Colored arrows are directions for mountain folding (**grey**; R-group in the front of plane, bold bond) and valley folding (**yellow**; R-group in the back of the plane, dashed bond). B) Folding symbols.

Fiber precursors are known as “**oligomers**”, and are small in size, which can be represented by single sheets containing 8 or 16 polypeptide chains. The spacing distance between polypeptide chains along the elongation axis is about 4.7 Å (i.e., 0.47 nm), and thus a one micrometer-long (1,000 nm) fiber could involve hydrogen-bonding 2,127 layers of polypeptides chains. Considering eight polypeptide chains in the elongation axis in a A4 paper sheet, it would require 265 printed and pleated paper sheets, connected to each other (about 56-meter-long), to represent a large mature fiber. Using the model with 16 chains per A4 paper sheet, half the amount of folded models would be required (28-meter-long). In a teaching class with several students each model can be connected using adhesive tape and/or aligned on a proper surface (floor, table, wall), aligned or hanging with threads. The dimensionality of a mature, long amyloid fiber can also be exemplified by using a thread as long as desired and the printed/folded amyloid origami model as a scale.

Topological representation of example amyloid structures are shown for aβ protein (**Fig. 4**) and α-Synuclein (**Fig. 5)**. Notice that the same protein (although with different chain length) can fold in different patterns. After pleating the paper in alternating folding creases using the bonds of adjacent chains as reference, the paper can be positioned upward, and the view from top representing a perspective of the topologic distribution of the polypeptide chain on the plane perpendicular to the fibril elongation axis (**Fig. 6**). Further topologic patterns can be obtained from high resolution amyloid structures solved and deposited in the RCSB, or from compilation deposited in databases such as in **Amyloid Atlas** (https://people.mbi.ucla.edu/sawaya/amyloidatlas/). Additional tools can be used to hold still the pleated paper representing the amyloid fold, such as elastic bands, transparent plastic cups or adhesive tape.

**Figure 4.**
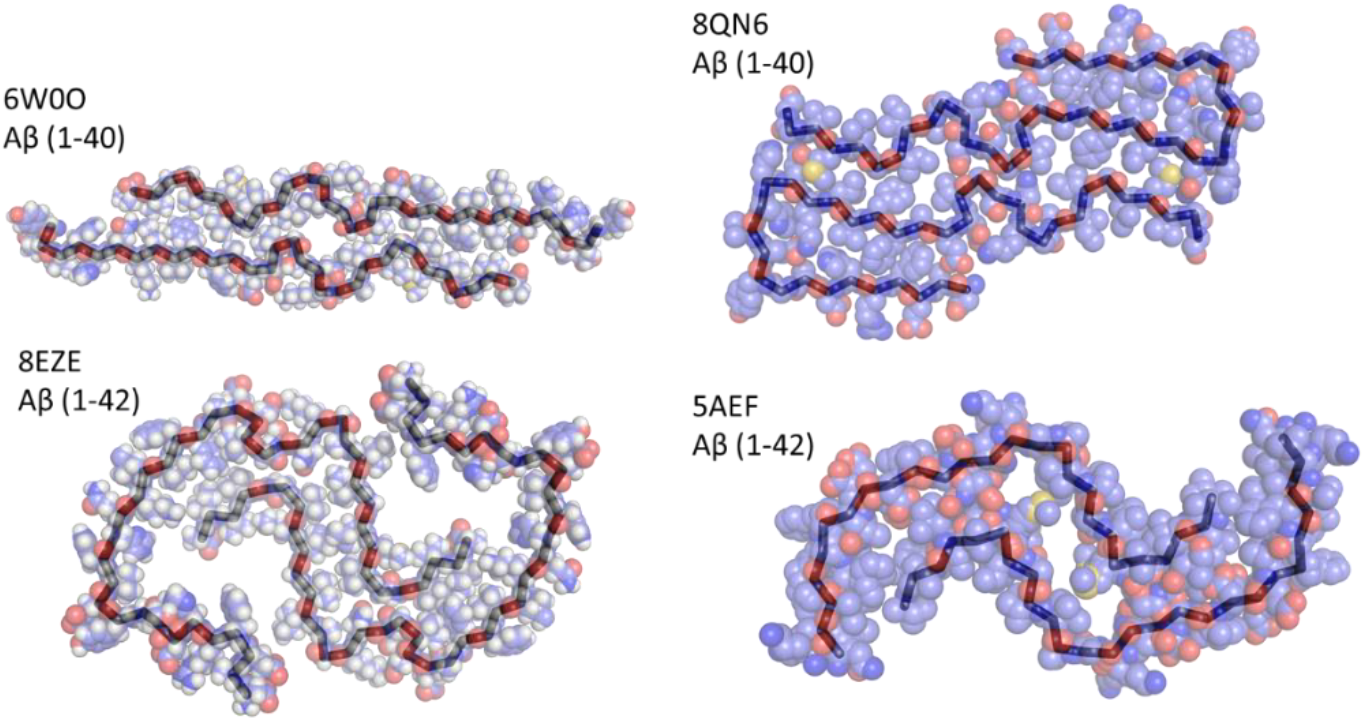
Topological representation of amyloid folding of amyloid-β (aβ). Codes correspond to PDB ID. Structures were created with PyMOL, using spheres transparency 40%, ribbon width 8, ribbon color black. Elements are represented as red for oxygen, blue for nitrogen, cyan for carbons, yellow for sulfur.

**Figure 5.**
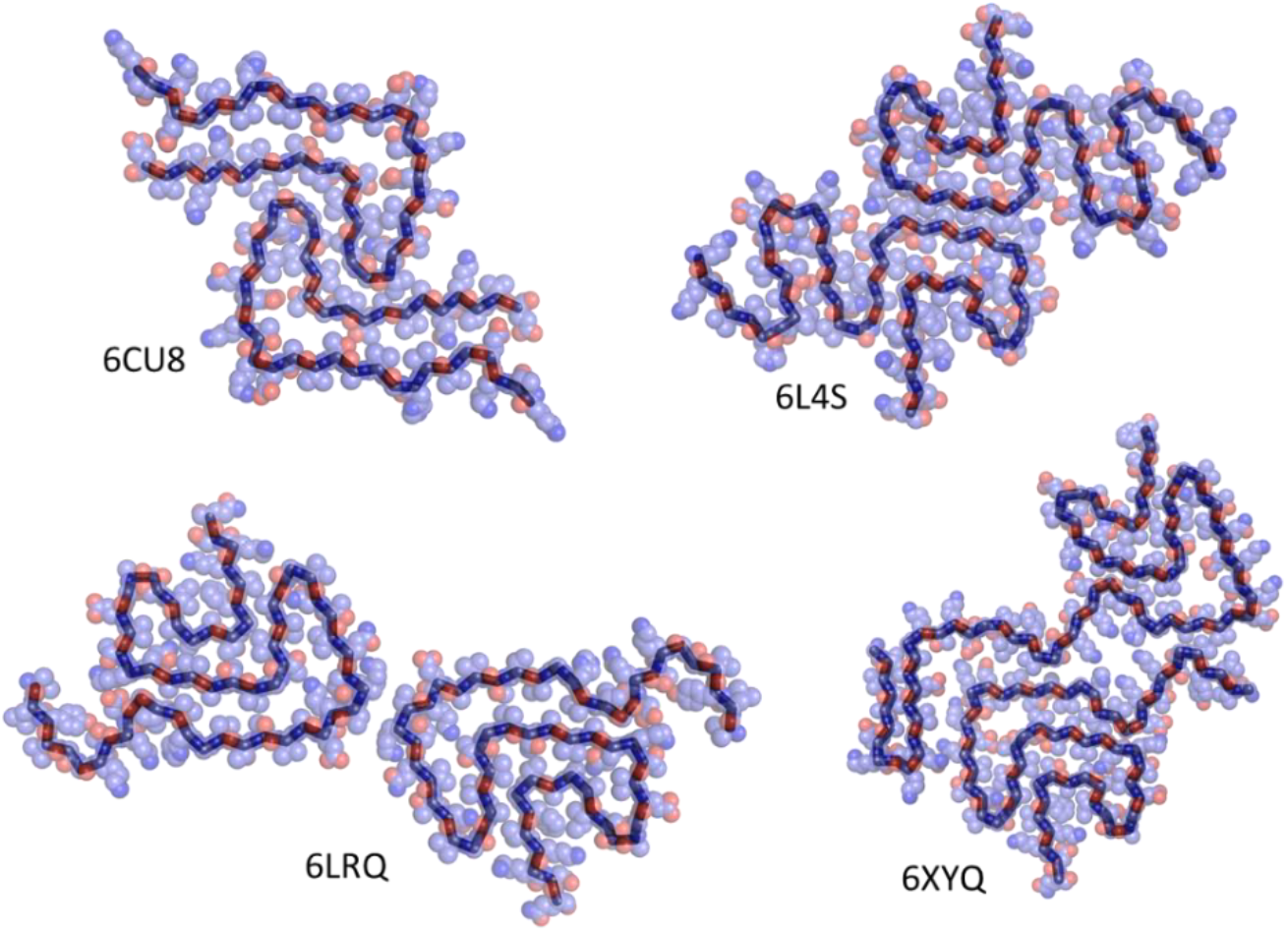
Topological representation of amyloid folding of α-synuclein. Codes correspond to PDB ID. Structures were created with PyMOL, using spheres transparency 40%, ribbon width 8, ribbon color black. Elements are represented as red for oxygen, blue for nitrogen, cyan for carbons, yellow for sulfur.

**Figure 6.**
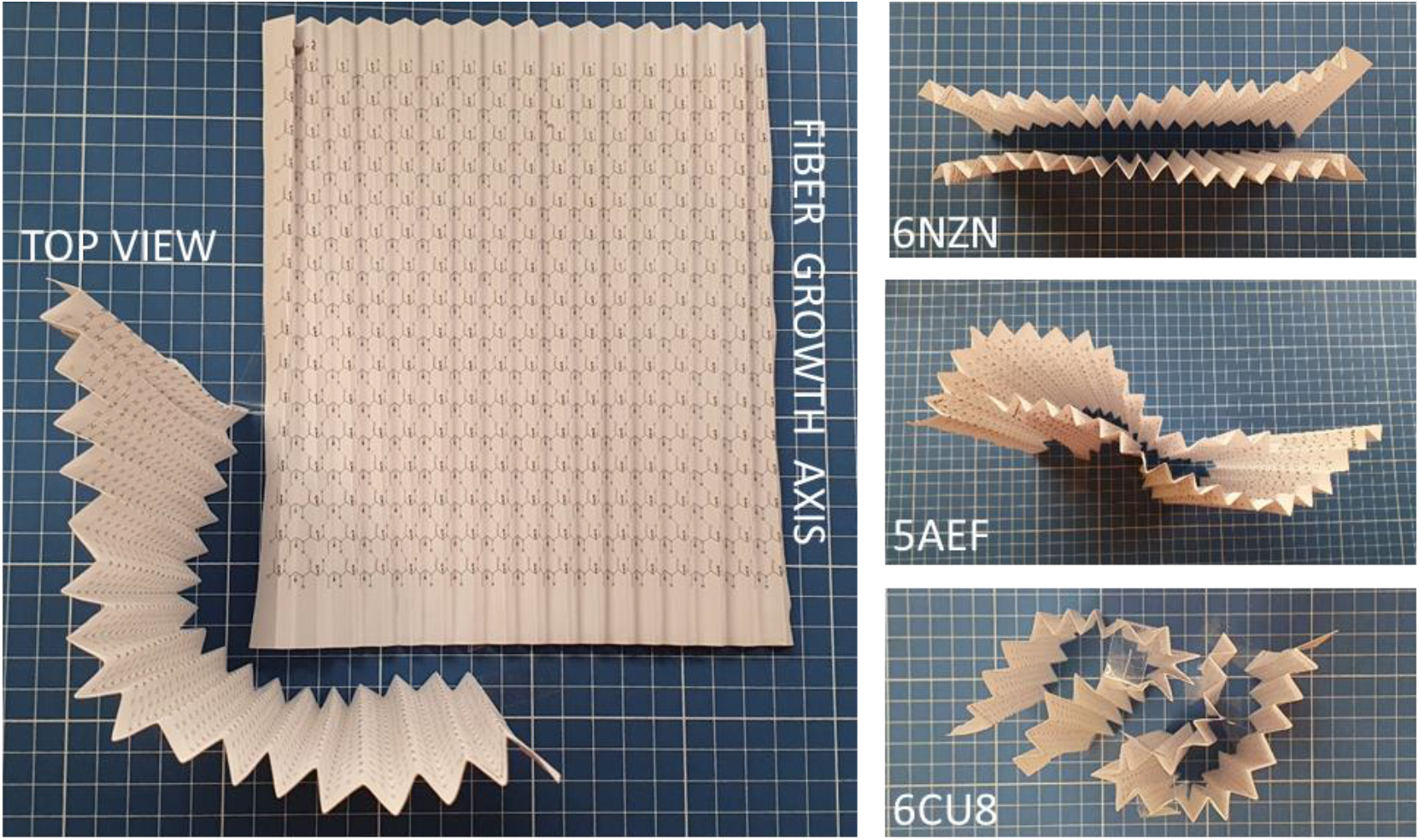
Origami representation of amyloid folding. Printed chains are folded in pleated pattern, alternating mountain / valley folds. Three arrangements are shown, representing glucagon (PDB ID 6NZN), Aβ (PDB ID 5AEF) and α-synuclein (PDB ID 6CU8) Not to scale between panels.

## 4. Discussion

Origami is a millenary art which uses paper for the representation of forms. Paper folding has also been used to represent and explain several physical and biological principles, such as the brain folding morphology (Mota & Herculano-Houzel 2015) and fractal (Balankin *et al*. 2010). While origami has long been used to assist understanding of globular protein folding, the nature of amyloid structures of protein fibers remains difficult to understand despite being described for close to one century since the seminal work of Prof. Astbury (Astbury 1931).

The experimental origami models presented here offer a tangible and intuitive representation of amyloid fibril architecture, translating molecular principles into a scalable and visual format. By folding printed sheets containing arrays of polypeptide chains—either in parallel or antiparallel orientations—key structural features of amyloids, such as hydrogen bonding along the elongation (Z) axis and side-chain interactions across stacking planes (X-Y), can be directly visualized and manipulated. This approach not only reinforces the geometric regularity of the cross-β pattern but also provides an educational bridge between abstract molecular structures and real-world spatial reasoning.

Because amyloids are most frequently associated with pathologies—such as Alzheimer’s disease (Aβ and tau), Parkinson’s disease (α-synuclein), type 2 diabetes mellitus, Creutzfeldt-Jakob disease, and various animal prion encephalopathies—their structures are often described in the context of “misfolding.” However, the term “misfolding” can be better associated to crumpled paper, which has a fractal pattern (Balankin *et al*. 2010), and more likely to be related to amorphous protein folding. Instead, amyloid structures are highly organized chemically and structurally, such as well-folded paper. In this context, our origami model uses pleated folds to represent the rigid amide bonds of the polypeptide backbone, with kinked points symbolizing the angles at α-carbon atoms where amino acid side chains emerge. The direction of the paper fold (mountain *vs* valley) represents the direction of the side-chain, which can promptly be associated with the complementary binding comprising the interaction between adjacent chains through polar and/or steric zippers. Moreover, the flexibility in the topological view allows understanding the chain flexibility and arrangements upon multiple protein patterns as allowed by chemical compatibility and geometric / stereochemical restriction. The flexibility of the paper, seen from top view, allows observation of the amyloid polymorph diversity, contrasting with the rigid structure of the β-sheet formed along the perpendicular paper direction representing the hydrogen bonding network.

This model can be extended to other fields, such as material science and understanding of topology of other polymers, such as polyamide. The origami representation of amyloid is not a substitute for a better, detailed visualization in sophisticated software. Instead, it targets an audience as an introductory tool, simple, cheap, accessible, and educational. Additionally, frequently in amyloid assembled between two chains they cross the topological planes, which can be better represented in paper folding by inclination of the models. In this context, the study of amyloid through the present paper representation aims to facilitate the perspective view of the amyloid folding arrangement instead of representing an accurate, high-resolution structure of amyloid which can be explored with more sophisticated computer tools. This origami tool is instead readily available to students at all levels with a simple office paper with the printed or, alternatively, hand-drawn polypeptide chains.

This origami model acts as a didactic bridge between complex structural data from state-of-the–art techniques such as cryoEM and ssNMR, and intuitive spatial understanding, enabling both researchers and students to physically explore amyloid polymorphism, chain orientation, and stacking patterns. It elegantly captures the duality inherent in amyloid structure—rigid β-sheets along the fibril elongation axis contrasted with flexible topological variation across stacking planes. Its low cost, reproducibility, and accessibility—requiring only free software, standard printers, and regular paper—make it an inclusive educational tool suitable for public high schools, undergraduates, and graduate students alike. Beyond amyloid research, this approach holds clear potential for transdisciplinary applications, extending its educational and conceptual value to diverse scientific fields such as health and material sciences.

The work of Sir D’Arcy Thompson find place here, not only by allowing modeling the “**form**” but also by demonstration of the “**growth**” process involved in amyloid elongation. A plethora of amyloid folding patterns can be represented, contemplated and studied, as well as the elongation mechanism of amyloid fibers can be addressed. The biology thus finds the mathematics, explaining “*tensions and conditions of equilibrium of the component molecules*”.

In folding paper to reveal the logic of proteins, we are reminded that complexity can arise from simplicity, and that form follows not only function, but also fundamental principles of geometry and interaction. This model, though modest, echoes the deeper order within biological matter—where art, physics, and life converge.

## 5. Conclusions

This foldable model represents the main features of amyloid structure, including the parallel or antiparallel alignment of chains, hydrogen bonding along the fibril axis, and steric or polar side-chain interactions between layers. By translating the cross-β architecture into a tactile and visual format, this model helps demystify amyloid assembly and enables hands-on exploration of fibril structure. Despite its simplicity, it is based on structural data and is a valuable tool not only for teaching the molecular basis of amyloid diseases, but also for communicating concepts of protein folding to a wider audience. In this way, it helps bridge the gap between atomically resolved structures and intuitive understanding, promoting access and stimulating interest in protein folding and aggregation and their impact on health and disease.

## Abbreviations

cryoEM: cryoelectron microscopy
ssNMR: solid-state Nuclear Magnetic Resonance
PDB ID: Protein Data Bank Identification (at https://www.rcsb.org/)

## Acknowledgments

Thanks for Laura, João and Alice Trambaioli Vieira Lima for unbiased (no prior knowledge about amyloid, protein, organic chemistry) testing the models, and to Prof. Mariana Quezado for revision and suggestions.

## Funding and additional information

This work was supported by Conselho Nacional de Desenvolvimento Científico e Tecnológico (CNPq; PQ/311784/2023-2, to LMTRL), Fundação de Amparo à Pesquisa do Estado do Rio de Janeiro Carlos Chagas Filho (FAPERJ; E-26/010.001434/2019-Tematico, E-26/210.195/2020 and SEI-260003/001207/2023 - APQ1 to LMTRL). The funding agencies had no role in the study design, data collection and analysis, or decision to publish or prepare of the manuscript.

### Conflict of Interest

The authors have no financial conflicts of interest with the contents of this article.

### Supporting Material

*ChemSketch* and PDF files of parallel / antiparallel chains, with 8 or 16 polypeptide chains.

## Footnotes

1 https://royalsocietypublishing.org/doi/pdf/10.1098/rsta.1934.0010 ; https://royalsocietypublishing.org/doi/10.1098/rsta.1934.0010

## Notes

### Competing Interest Statement

The authors have declared no competing interest.

## REFERENCES

Astbury WT 1931 X-ray studies of the structure of hair, wool, and related fibres.-I. General. Philosophical Transactions of the Royal Society of London. Series A, Containing Papers of a Mathematical or Physical Character. (doi:10.1098/rsta.1932.0003)

Astbury WT 1933 X-Ray studies of the structure of hair, wool, and related fibres. II.-the molecular structure and elastic properties of hair keratin. Philosophical Transactions of the Royal Society of London. Series A, Containing Papers of a Mathematical or Physical Character. (doi:10.1098/rsta.1934.0010)

Azulay H, Pellach Leshem M & Qvit N 2020 An Approach to comparing protein structures and origami models - Part 2. Multi-domain proteins. Biochimica Et Biophysica Acta. Biomembranes 1862 183411. (doi:10.1016/j.bbamem.2020.183411)

Azulay H, Lutaty A & Qvit N 2022 How Similar Are Proteins and Origami? Biomolecules 12 622. (doi:10.3390/biom12050622)

Balankin AS, Ochoa DS, Miguel IA, Ortiz JP & Cruz MÁM 2010 Fractal topology of hand-crumpled paper. Physical Review E 81 061126. (doi:10.1103/PhysRevE.81.061126)

Buxbaum JN, Dispenzieri A, Eisenberg DS, Fändrich M, Merlini G, Saraiva MJM, Sekijima Y & Westermark P 2022 Amyloid nomenclature 2022: update, novel proteins, and recommendations by the International Society of Amyloidosis (ISA) Nomenclature Committee. Amyloid.

Chiti F, Webster P, Taddei N, Clark A, Stefani M, Ramponi G & Dobson CM 1999 Designing conditions for in vitro formation of amyloid protofilaments and fibrils. Proceedings of the National Academy of Sciences 96 3590–3594. (doi:10.1073/pnas.96.7.3590)

De Baere I, Liu L, Moens L, Van Beeumen J, Gielens C, Richelle J, Trotman C, Finch J, Gerstein M & Perutz M 1992 Polar zipper sequence in the high-affinity hemoglobin of Ascaris suum: amino acid sequence and structural interpretation. Proceedings of the National Academy of Sciences 89 4638–4642. (doi:10.1073/pnas.89.10.4638)

Demaine ED & O’Rourke J 2007 Geometric Folding Algorithms: Linkages, Origami, Polyhedra. Cambridge: Cambridge University Press.

Garratt RC, Beltramini LM & Abel LD dos S 2015 [Kit didático: construindo modelos topológicos de proteínas].

Ivanova MI, Sievers SA, Sawaya MR, Wall JS & Eisenberg D 2009 Molecular basis for insulin fibril assembly. Proceedings of the National Academy of Sciences 106 18990–18995. (doi:10.1073/pnas.0910080106)

Kendrew JC 1961 The three-dimensional structure of a protein molecule. Scientific American 205 96–110. (doi:10.1038/scientificamerican1261-96)

Kerwin SM 2019 Flexible and modular 3D-printed peptide models. Biochemistry and Molecular Biology Education: A Bimonthly Publication of the International Union of Biochemistry and Molecular Biology 47 432–437. (doi:10.1002/bmb.21250)

Landschulz WH, Johnson PF & McKnight SL 1988 The Leucine Zipper: A Hypothetical Structure Common to a New Class of DNA Binding Proteins. Science 240 1759–1764. (doi:10.1126/science.3289117)

Levinthal C 1969 How to Fold Graciously. In Mossbauer Spectroscopy in Biological Systems: Proceedings of a Meeting Held at Allerton House, Monticello, Illinois, pp 22–24. Allerton House, Monticello, Illinois.: University of Illinois Press.

Meyer SC 2015 3D Printing of Protein Models in an Undergraduate Laboratory: Leucine Zippers. Journal of Chemical Education 92 2120–2125. (doi:10.1021/acs.jchemed.5b00207)

“Molecular Origami: Build 3D models of PDB Structures.”

“Molecular Origami: Build 3D Paper Models of Protein Domains.”

Mota B & Herculano-Houzel S 2015 BRAIN STRUCTURE. Cortical folding scales universally with surface area and thickness, not number of neurons. Science (New York, N.Y.) 349 74–77. (doi:10.1126/science.aaa9101)

Perutz M 1994 Polar zippers: their role in human disease. Protein Science: A Publication of the Protein Society 3 1629–1637. (doi:10.1002/pro.5560031002)

Perutz MF, Rossmann MG, Cullis AF, Muirhead H, Will G & North ACT 1960 Structure of Hæmoglobin: A Three-Dimensional Fourier Synthesis at 5.5-Å. Resolution, Obtained by X-Ray Analysis. Nature 185 416–422. (doi:10.1038/185416a0)

Perutz MF, Staden R, Moens L & De Baere I 1993 Polar zippers. Current Biology 3 249–253. (doi:10.1016/0960-9822(93)90174-M)

Popa I & Saitis F 2022 Using Magnets and Flexible 3D-Printed Structures to Illustrate Protein (Un)folding. Journal of Chemical Education 99 3074–3082. (doi:10.1021/acs.jchemed.2c00231)

Reißer S, Prock S, Heinzmann H & Ulrich AS 2018 Protein ORIGAMI: A program for the creation of 3D paper models of folded peptides. Biochemistry and Molecular Biology Education: A Bimonthly Publication of the International Union of Biochemistry and Molecular Biology 46 403–409. (doi:10.1002/bmb.21132)

Sawaya MR, Hughes MP, Rodriguez JA, Riek R & Eisenberg DS 2021 The expanding amyloid family: Structure, stability, function, and pathogenesis. Cell 184 4857–4873. (doi:10.1016/j.cell.2021.08.013)

Thompson DW, Thompson K & Biology 1942 On Growth and Form: The Complete Revised Edition. New York: Dover Publications.

Watson JD & Crick FHC 1953 Molecular Structure of Nucleic Acids: A Structure for Deoxyribose Nucleic Acid. Nature 171 737–738. (doi:10.1038/171737a0)

