## Supporting Material for "Amyloid Folding: an Origami-Based Approach"

[1] Laboratório de Biotecnologia Farmacêutica – pbiotech, Faculdade de Farmácia, Universidade Federal do Rio de Janeiro – UFRJ.
[2] Programa de Pós-Graduação em Ciências Farmacêuticas, Faculdade de Farmácia, Universidade Federal do Rio de Janeiro, Rio de Janeiro, RJ, 21941-902, Brazil.
[3] Programa de Pós-Graduação em Química Biológica, Universidade Federal do Rio de Janeiro, Rio de Janeiro, RJ, 21941-902, Brazil.
[4] Programa de Pós-Graduação em Nutrição, Universidade Federal do Rio de Janeiro, Rio de Janeiro, RJ, 21941-902, Brazil.

*Running Title:* Amyloid Origami

*To whom correspondence should be addressed*: Luís Maurício T. R. Lima – Faculdade de Farmácia, Universidade Federal do Rio de Janeiro – UFRJ, CCS, Bss24, Ilha do Fundão, 21941-902, Rio de Janeiro, RJ, Brazil. Phone/Fax: (+55-21) 3938-6639.

**Author information**

This is a pdf version of the associated *ChemSketch* file, containing four groups distributed in the following eight pages:

**Group 1**
- 8 parallel chains, front
- 8 parallel chains, back (flipped)

**Group 2**
- 8 anti-parallel chains, front
- 8 anti-parallel chains, back (flipped)

**Group 3**
- 16 parallel chains, front
- 16 parallel chains, back (flipped)

**Group 4**
- 16 anti-parallel chains, front
- 16 anti-parallel chains, back (flipped)

Each group should be printed one in each side of the paper (front & back pair) in A4 format.

Diagrams can be printed in varying paper sizes, such as office, letter, A4, A3, A2 or other according to user preferences and availability, preferably from the associated *ChemSketch* file using the freely available ChemSketch software (https://www.acdlabs.com/resources/free-chemistry-software-apps/chemsketch-freeware/).

After printing out the desired models, follow the folding instructions as depicted in the manuscript, summarized below:

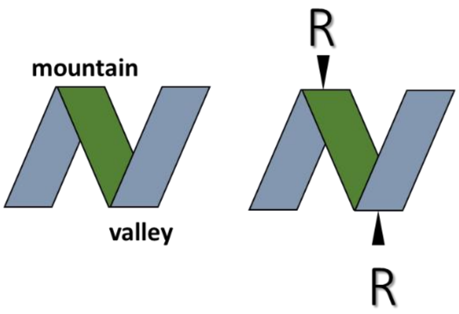


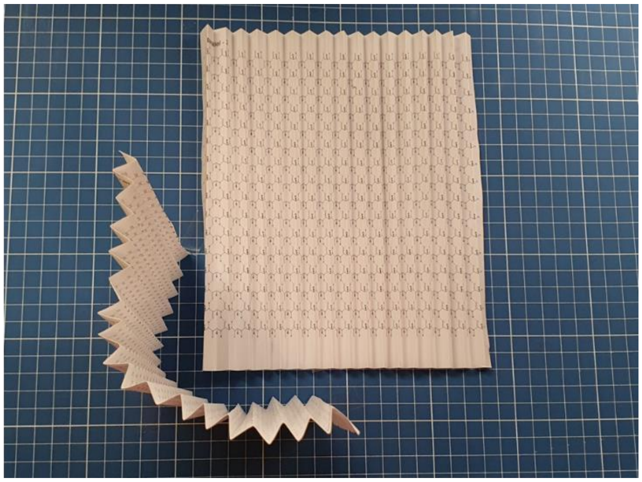

Amyloid folding topologies can be found in the RCSB from high-resolution structures, and/or from the compilation database **Amyloid Atlas** (https://people.mbi.ucla.edu/sawaya/amyloidatlas/).

parallel - 1

**Amyloid Origami**

*Luis Mauricio Trambaioli da Rocha e Lima*

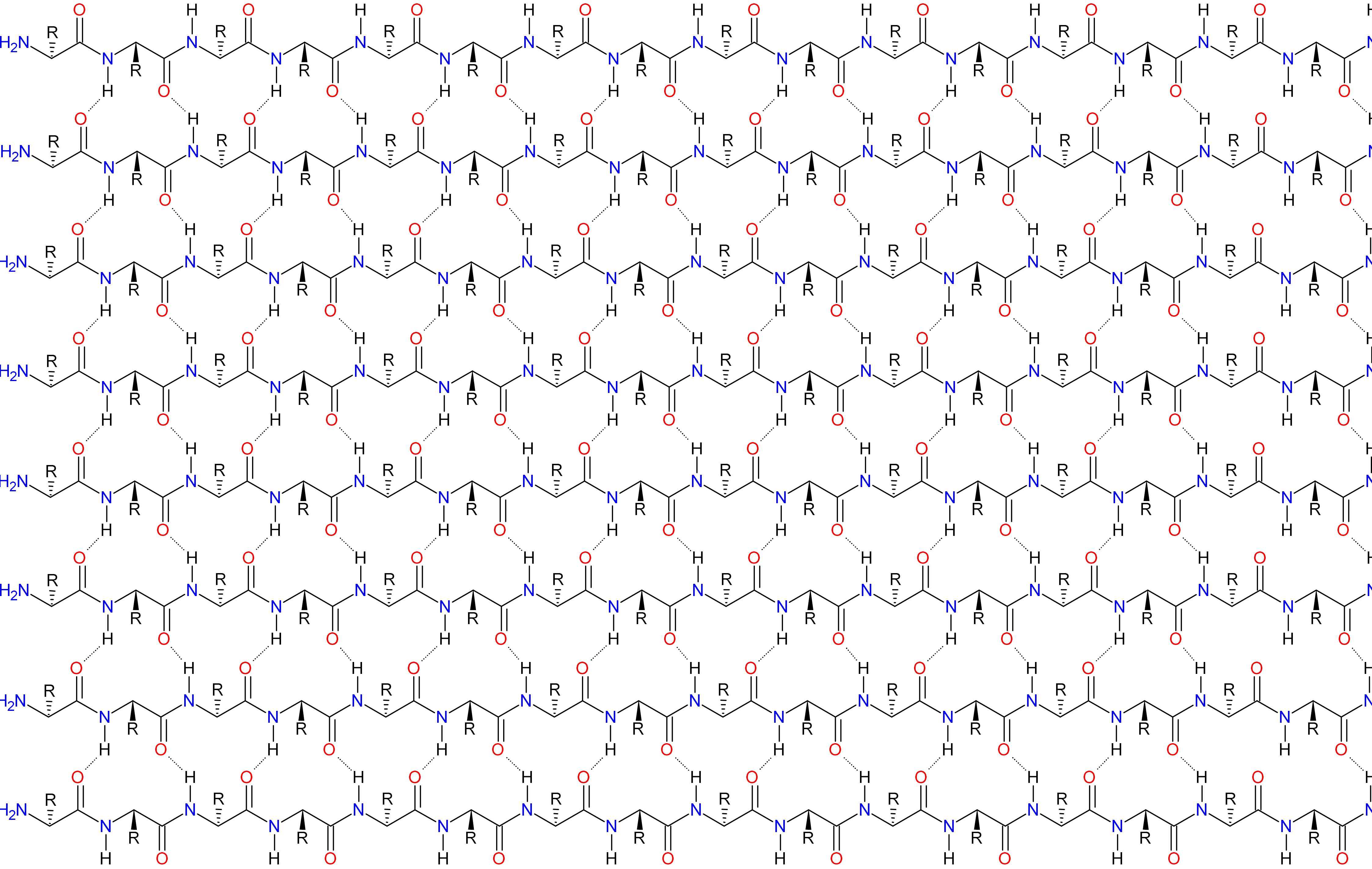

parallel - 2

**Amyloid Origami**

*Luis Mauricio Trambaioli da Rocha e Lima*

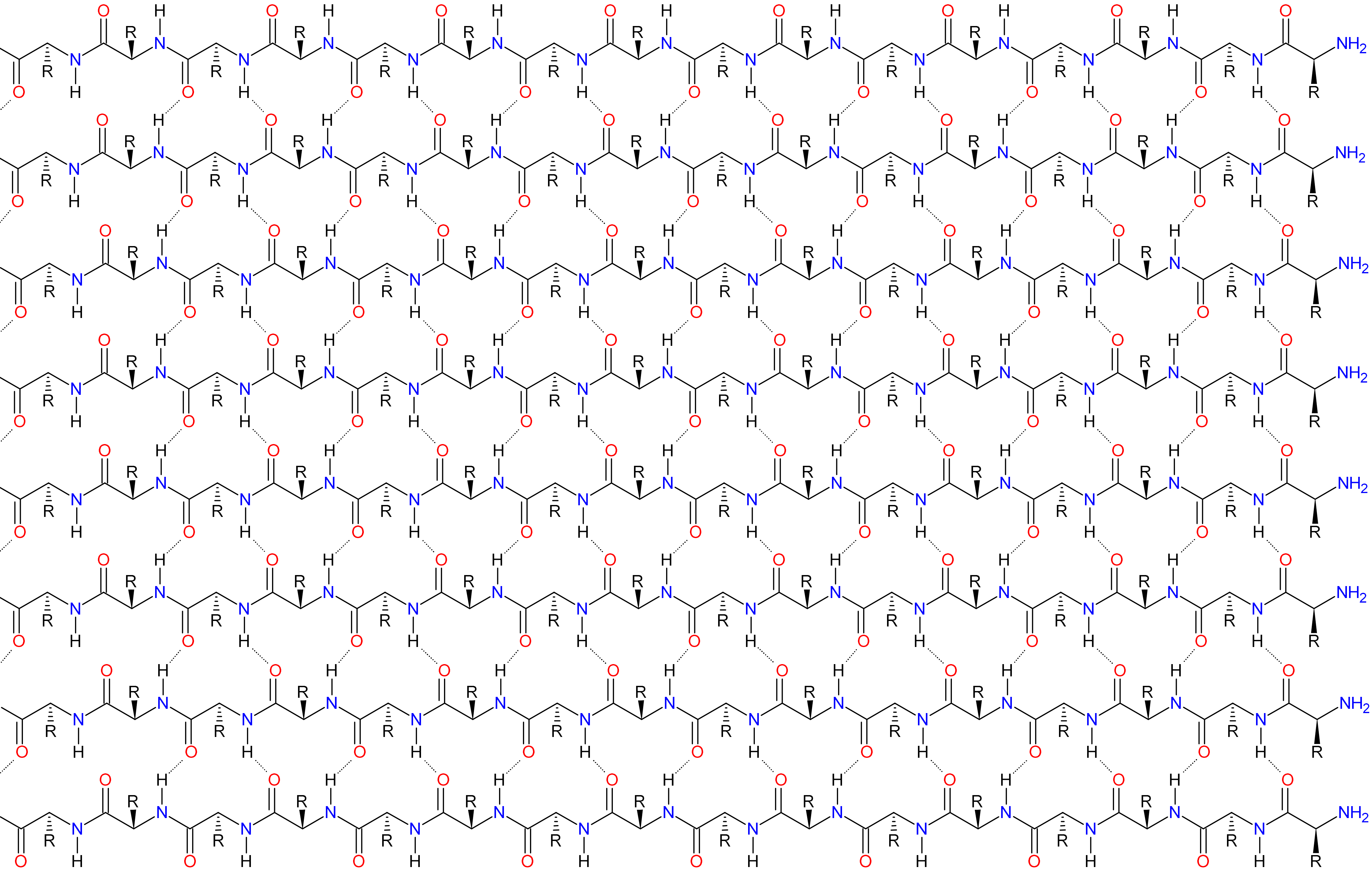

antiparallel - 1

**Amyloid Origami**

*Luis Mauricio Trambaioli da Rocha e Lima*

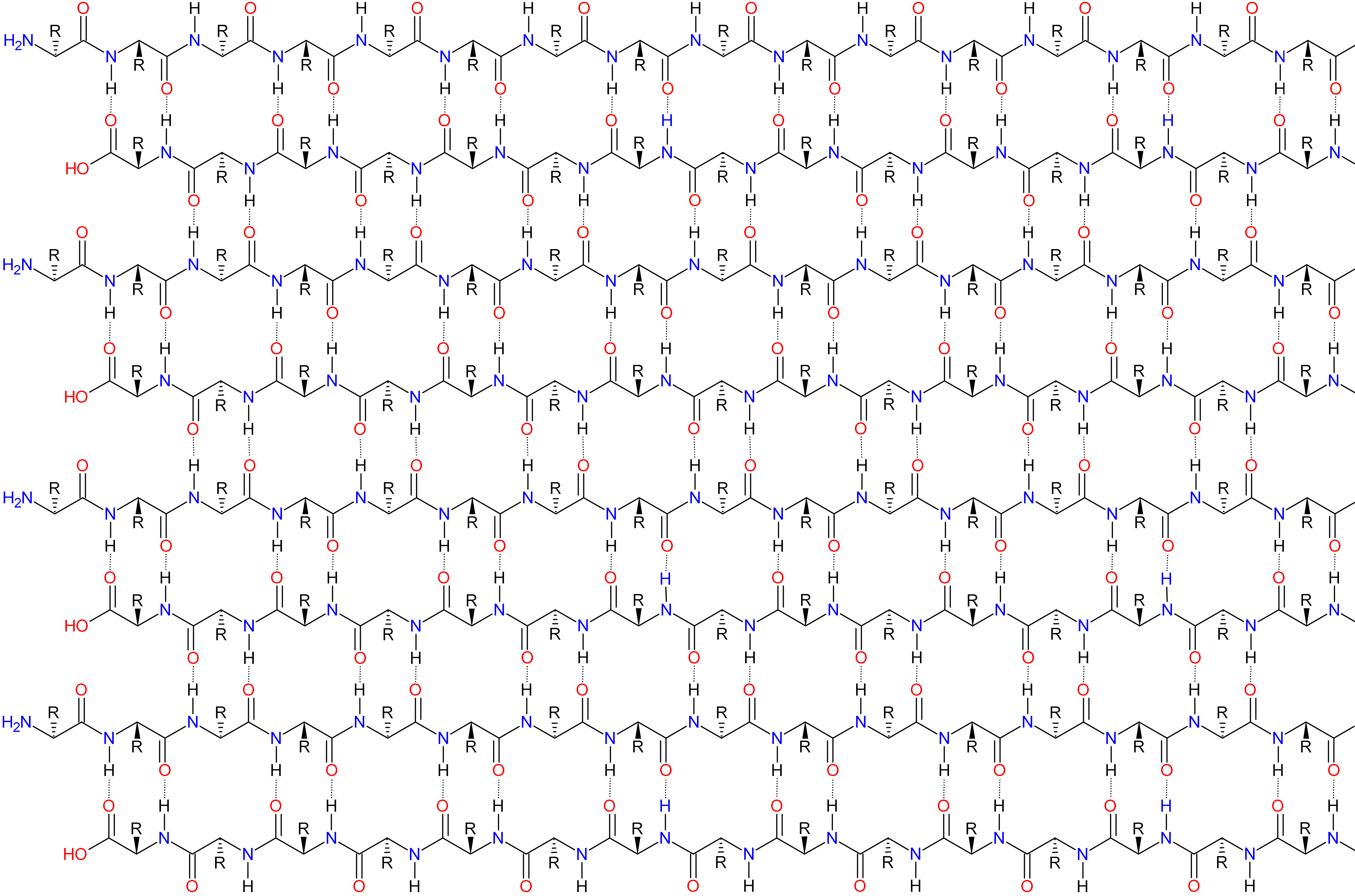

antiparallel - 2

**Amyloid Origami**

*Luis Mauricio Trambaioli da Rocha e Lima*

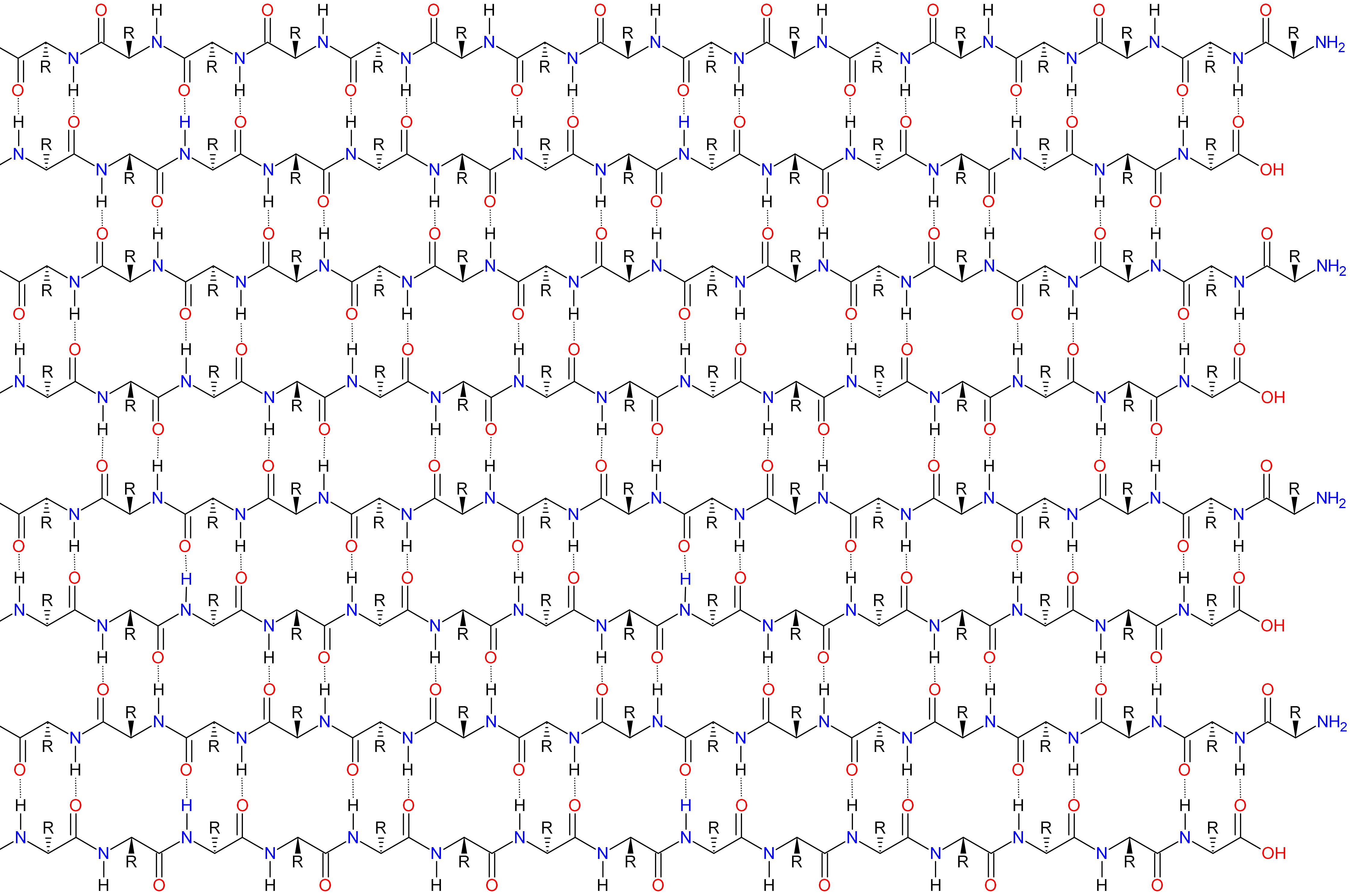

parallel - 1

### Amyloid Origami

*Luis Mauricio Trambaioli da Rocha e Lima*

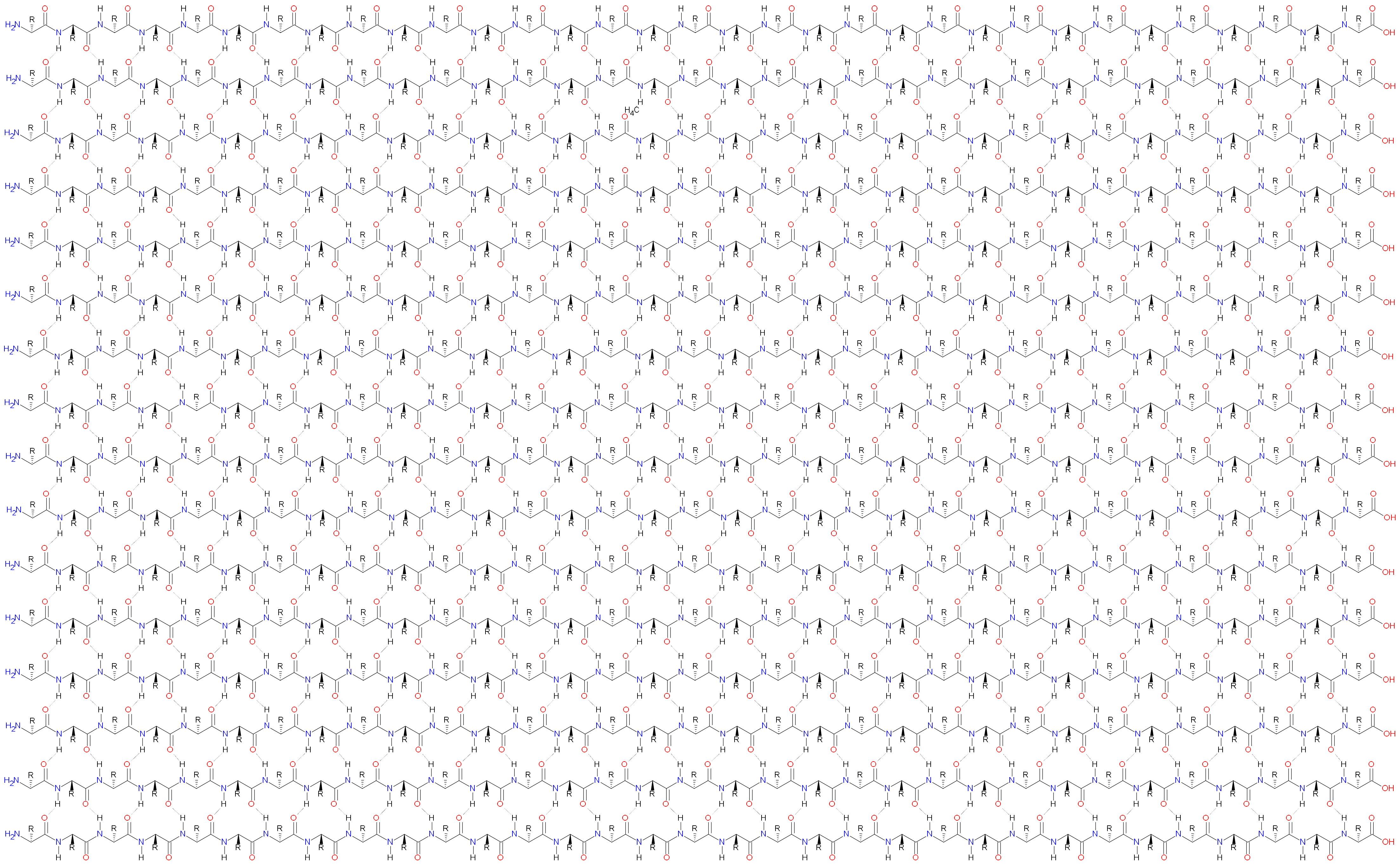

### Amyloid Origami

*Luis Mauricio Trambaioli da Rocha e Lima*

parallel - 2

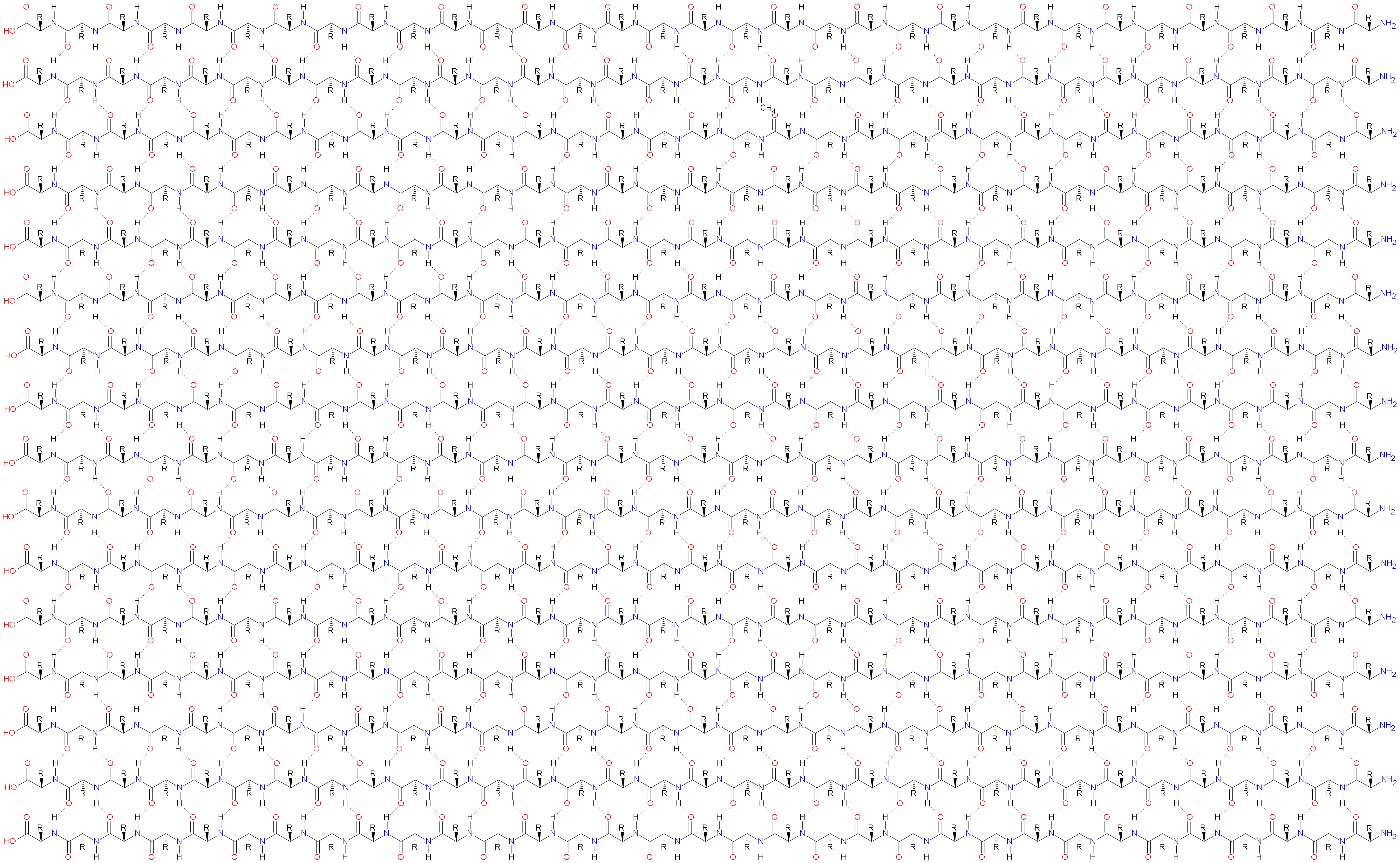

antiparallel - 1

**Amyloid Origami**

*Luis Mauricio Trambaioli da Rocha e Lima*

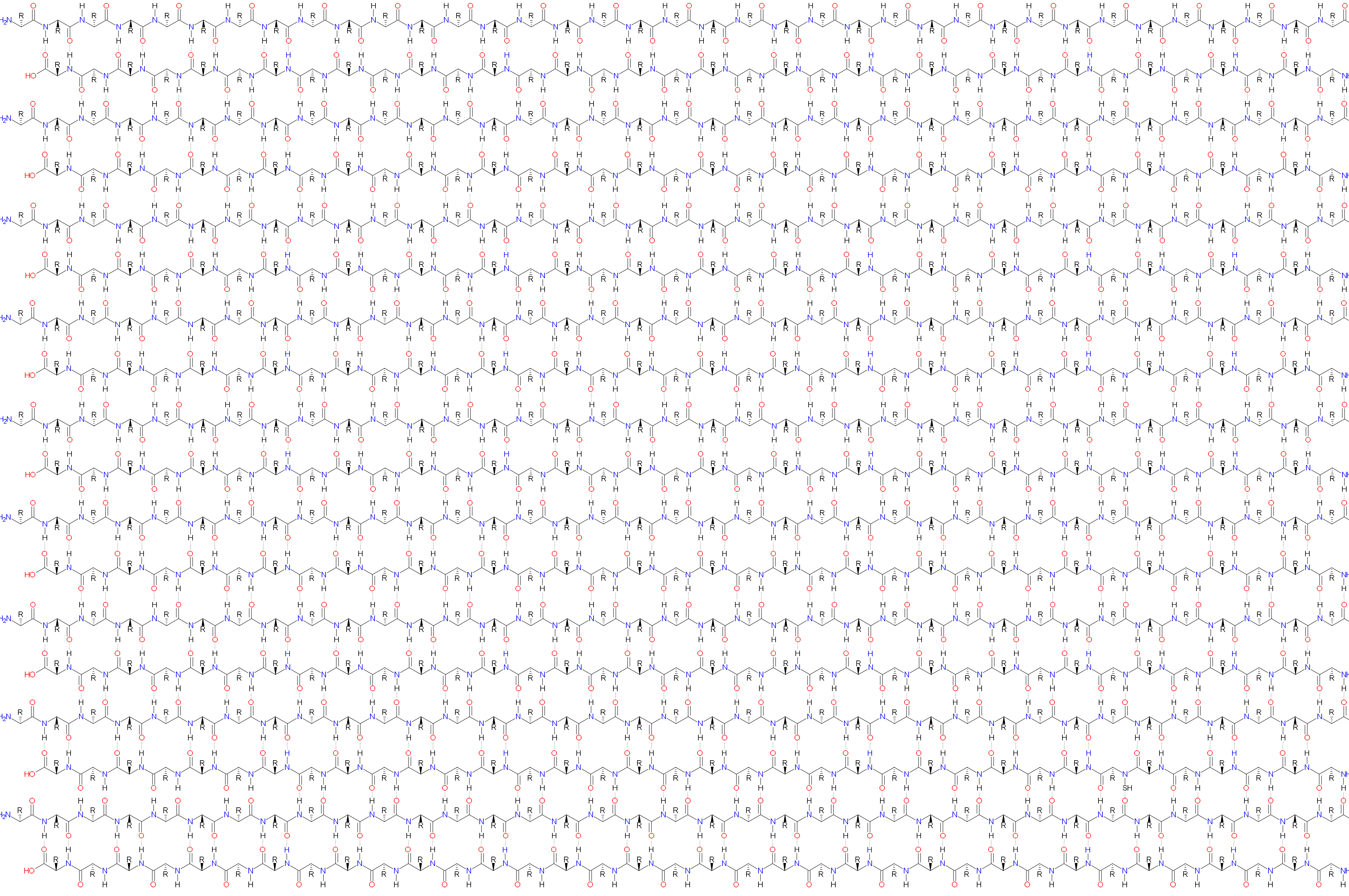

### Amyloid Origami

*Luis Mauricio Trambaioli da Rocha e Lima*

antiparallel - 2

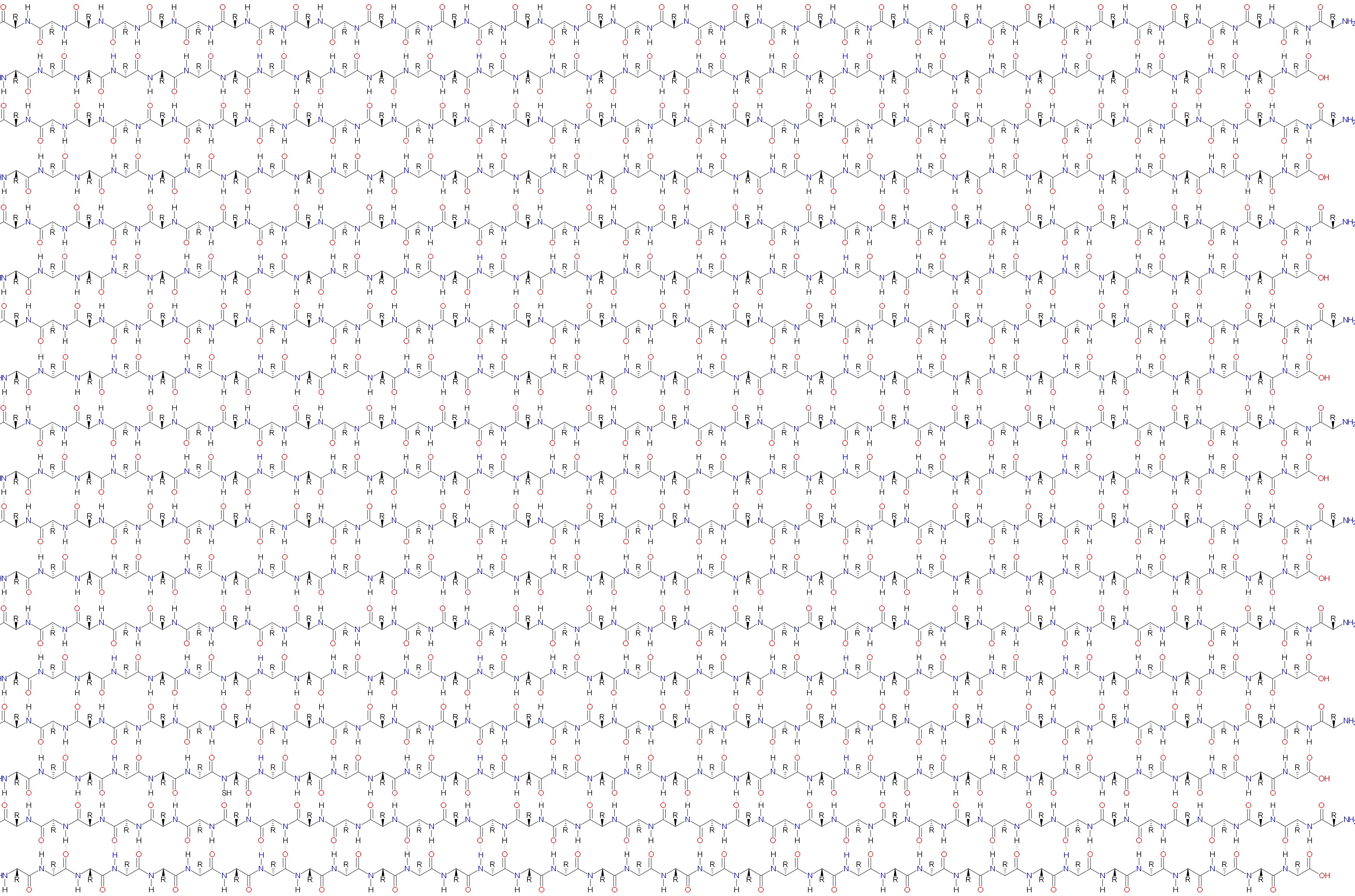
